# A novel fluorescently-labeled long-chain fatty acid analog for the study of fatty acid metabolism in cultured cells

**DOI:** 10.1101/2022.12.06.519076

**Authors:** Yasuhiro Hara, Ken-ichi Hirano

## Abstract

^123^I-15-(p-iodophenyl)-(*R,S*)-methyl pentadecanoic acid (BMIPP) is a long-chain fatty acid (LCFA) analog developed to examine myocardial LCFA metabolism and has been used as a tracer for nuclear cardiology. However, its use is limited because of the specialized features of cardiac scintigraphy. In this study, a novel BMIPP-based probe was utilized, in which iodine-123 was replaced with a fluorescent compound, to extend the use of ^123^I-BMIPP to a wider variety of cells *ex vivo*. To confirm that this fluorescent LCFA analog (fluorescent BMPP) was imported into cells, fluorescence-activated cell sorting (FACS) analysis and fluorescent cell imaging were performed using cultured cells. The analysis showed that the import of fluorescent BMPP into the cells occurred in a concentration-dependent manner. This import into cells was inhibited by Sulfosuccinimidyl Oleate in a dose-dependent manner, which is an inhibitor of CD36, a well-known LCFA transporter, suggesting that fluorescent BMPP could be imported into cells via the same pathway as LCFA. FACS and cell imaging intensities of the cells importing fluorescent BMPP were attenuated after incubation in the non-Alexa680-BMPP medium. These results suggest that fluorescent BMPP can be transported into and from cells, reflecting the metabolism of LCFA. Fluorescently-labeled BMPP has the potential to be used as a probe for studying LCFA metabolism in various cells.

## 1. Introduction

We aimed to develop a useful *ex vivo* probe capable of examining intracellular fatty acid (FA) metabolism in different types of cells, and to eventually develop a diagnostic system capable of detecting intracellular triglyceride (TG) hydrolysis ability in heart diseases *ex vivo*. One of our target diseases is triglyceride deposit cardiomyovasculopathy (TGCV), which is caused by a genetic or acquired deficiency of adipose triglyceride lipase, a rate-limiting enzyme for intracellular TG hydrolysis, resulting in excessive accumulation of TG in cardiomyocytes and the coronary artery [1-3]. We previously established the diagnostic criteria for TGCV and demonstrated the increasing importance of ^123^I-15-(p-iodophenyl)-(*R,S*)-methyl pentadecanoic acid (BMIPP) cardiac scintigraphy using single-photon emission computed tomography (SPECT) imaging [3-5]. The long-chain fatty acid (LCFA) analog ^123^I-BMIPP has been used in nuclear cardiology to study FA metabolism in different types of heart disease [6-9], including TGCV.

However, the use of iodine-123 in scintigraphy with SPECT equipment has disadvantages such as the need for appropriate facilities. Therefore, we focused on developing a novel fluorescently-labeled BMIPP-basic structure without using a radioisotope. In a previous study, we synthesized a BMIPP-based fluorescent LCFA analog, in which iodine-123 was replaced with Alexa Fluor 680 [10]. In this study, we investigated the characteristics of fluorescently-labeled BMIPP skeleton to detect FA metabolism in cultured cells. We demonstrated that the fluorescently-labeled BMIPP skeleton was successfully transported into and from the cells.

## 2. Materials and Methods

### 2.1 Cell culture

HeLa and H9C2 cells were maintained in high-glucose Dulbecco’s modified Eagle’s medium (DMEM) (Nacalai Tesque, Kyoto, Japan) supplemented with 10% fetal bovine serum (FBS) (Gibco / Thermo Fisher Scientific, Waltham, MA, USA) and 100 μg/mL streptomycin at 37 °C with 5% CO_2_.

### 2.2 Treatment of cells with Fluorescently-labeled BMPP

Fluorescently-labeled BMPPs (fluorescent BMPP) in phosphate buffered saline (PBS) were kindly gifted by Professors Kenji Monde and Takashi Jin, and were prepared from amino-BMPP as described in a previous report [10]. All experiments were performed at least twice. Fresh non-FBS DMEM containing fluorescent BMPP (concentrations indicated in figure legends) was added to cells grown to 60–80% confluence in 6-well culture plates or 35-mm culture dishes and incubated for 30 min under normal culture conditions (see Section 2.1). The cells were then subjected to fluorescence-activated cell sorting (FACS) analysis or fluorescent cell imaging.

### 2.3 FACS analysis of the import of fluorescent BMPP into cells

After treatment with fluorescent BMPP, cells were washed with PBS, detached by incubation in trypsin/EDTA solution (Nacalai Tesque Inc.) for 4 min at 37 °C, and centrifuged at 1000 × *g* for 2 min. The resulting cell pellets were resuspended in 500 μL of PBS or DMEM (-FBS) and filtered using a 35-μm cell strainer (BD Biosciences, San Jose, CA, USA) into single-cell suspensions for FACS analysis.

The fluorescence intensities of the recovered cells were analyzed using a MacsQuant Analyzer flow cytometer (Miltenyi Biotec, Bergisch Gladbach, Germany) according to the manufacturer’s instructions. Briefly, the flow cytometer was set to measure side scatter (SSC) and forward scatter (FSC), and FL2 (ex/em, 488/525 nm) for fluorescein fluorescence, and channel FL6 (ex/em, 635/655–730 nm) for Alexa680 fluorescence. The SSC and FSC amplifications were adjusted using non-fluorescent control cells recovered using the same protocol prior to measuring the fluorescent signals to determine a gating region including the major cell populations, except for debris. Fluorescein and Alexa680 fluorescence parameters were adjusted using control cells prior to sample measurement to obtain suitable peak positions. Finally, 10^4^ cells (in 200–400 μL of PBS or DMEM) were measured for the single-color fluorescence analysis of either fluorescein or Alexa680.

### 2.4 Imaging of the import of fluorescent BMPP into cells

Cells were grown in 35-mm glass bottom culture dishes to 60–80% confluence and then treated with fluorescent BMPP as described above. Hoechst 33342 was also added to the medium (5 μg/mL) for nuclear staining. After washing twice with PBS at the indicated time points, cells were fixed with 1:10 diluted formalin in PBS for 10 min, washed three times with PBS, and treated with the appropriate amount of PBS. Fluorescence images were acquired using a BZ-X700 microscope (Keyence, Osaka, Japan). The filters (ex/em, 470/525 nm), (ex/em, 620/700 nm), and (ex/em, 360/460 nm) were used to detect fluorescein, Alexa680, and Hoechst 33342, respectively.

### 2.5 Treatment of cells with Sulfosuccinimidyl Oleate (SSO)

Fresh DMEM (-FBS) containing SSO (Toronto Research Chemicals, North York, Canada) (concentrations indicated in figure legends) was added to the cells in 6-well culture plates or 35-mm glass bottom culture dishes and pre-treated for 30 min under normal culture conditions (see Section 2.1) prior to treatment with fluorescein-labeled BMPP (fluorescein-BMPP). The medium was then changed to DMEM (-FBS) containing SSO (the same concentrations as pre-treatment) and 0.05 μM fluorescein-BMPP. FACS analysis and cell imaging for fluorescein-BMPP-import were performed as described above.

### 2.6 Fluorescein-BMPP-import and export assay using FACS analysis

Cells in 6-well culture plates were treated with 0.075 μM fluorescein-BMPP, as described above. The export assay was performed as follows. After incubation for 30 min for fluorescein-BMPP import, cells were analyzed by FACS immediately after import. On the other hand, the export assay was performed using remaining import wells as follows. Cells were washed twice with DMEM to remove fluorescein-BMPP from the extracellular medium, and were added with fresh DMEM(+ FBA), and then incubated at 37 °C under normal culture conditions (see Section 2.1) for time periods as indicated in Figure. After incubation, the cells were analyzed by FACS. Cell detachment and FACS analyses were performed using the methods described above.

### 2.7 Fluorescein-BMPP-import and export assay using Cell imaging

Cells in 35-mm glass-bottom culture dishes were treated with 0.075 μM fluorescein-BMPP, as described above. An export assay using cell imaging was performed as follows. After incubation for 30 min for fluorescein-BMPP import, cells were fluorescently imaged immediately after import. On the other hand, the export assay was performed using remaining import dishes as follows. Cells were washed twice with DMEM to remove fluorescein-BMPP from the extracellular medium, and were added with fresh DMEM(+ FBA), and then incubated at 37 °C under normal culture conditions (see Section 2.1) for time periods as indicated in Figure. After incubation, cells were fluorescently imaged. Fluorescence imaging was performed as described above.

## 3. Results

### 3.1 Fluorescently-labeled basic structure of BMIPP was imported into cultured cells in dose-dependent manner

Our goal was to utilize a fluorescently-labeled BMIPP basic skeleton (defined fluorescent BMPP) as a potent probe for examining FA metabolism in cells *ex vivo*. Thus, we investigated the import of fluorescent BMPP into cells using FACS analysis and fluorescent cell imaging (Fig. 1 and 2). The results clearly showed that both Alexa680- and fluorescein-labeled BMPP were imported into HeLa and H9C2 cells in a BMPP concentration-dependent manner. In contrast, cells treated with non-reactive, hydrolyzed fluorescein isothiocyanate (FITC) (FITC control) and Alexa Fluor 680 (Alexa Fluor 680 control) showed the same intensity pattern as the non-added control (Fig.1 and 2). These results confirmed that the dose-dependent import of fluorescent BMPP into cells depended on the BMIPP basic skeleton as an LCFA analog.

**Figure 1.**
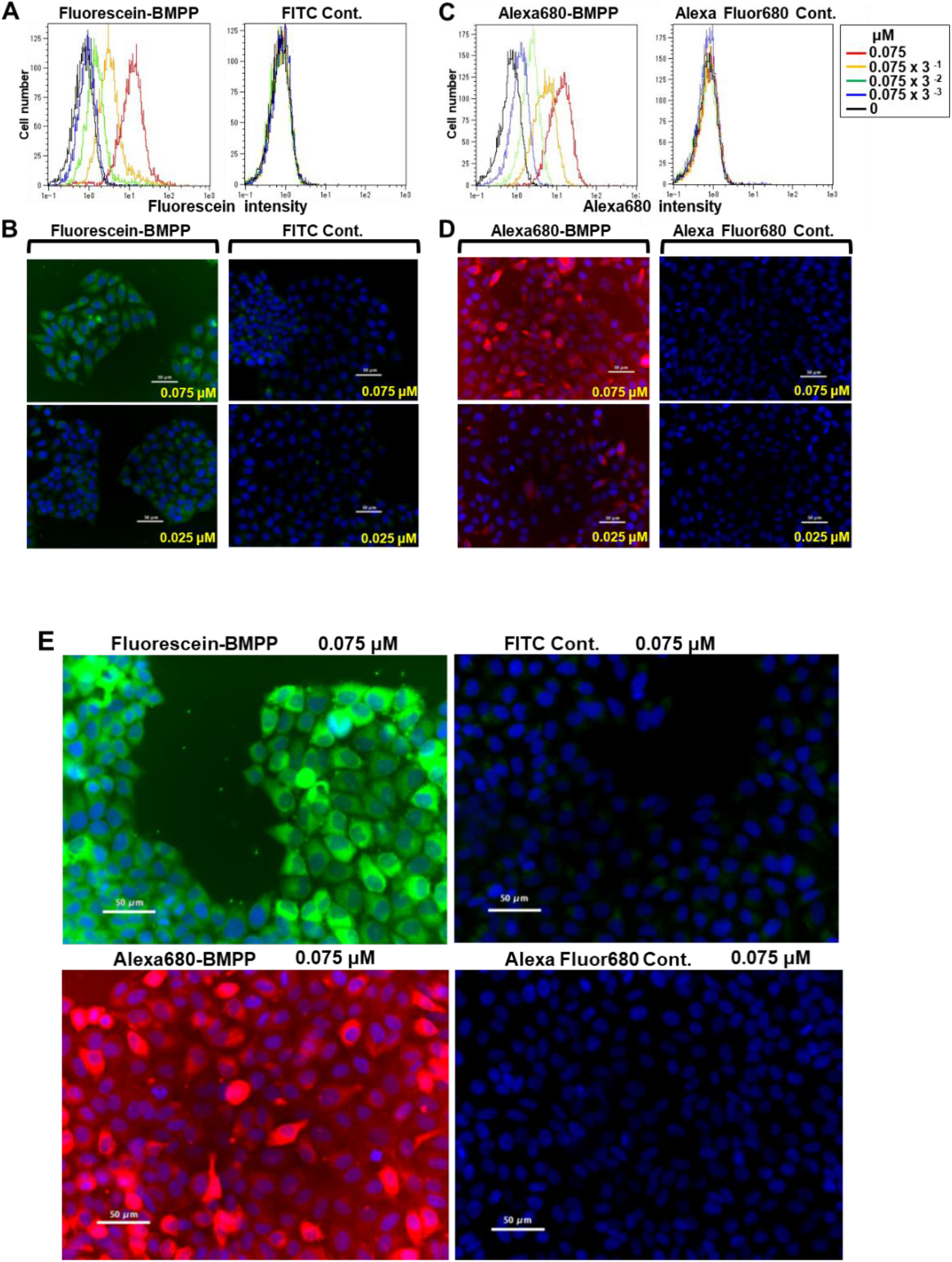
Dose-dependent import of fluorescent BMPP into HeLa cells. (A) FACS analysis of fluorescein-BMPP-treated HeLa cells. Cells were treated with increasing doses of fluorescein-BMPP (left) or non-reactive, hydrolyzed FITC (FITC Cont.) (right) (0.075 × 3^-3^ to 0.075 μM). X-axis of each graph represents fluorescein intensity. (B) Cell imaging of fluorescein-BMPP-treated HeLa cells. Cells were treated with fluorescein-BMPP (left) or FITC Cont.(right) (0.025 and 0.075 μM). Scale bar, 50 μm. (C) FACS analysis of Alexa680-BMPP-treated HeLa cells. Cells were treated with increasing doses of Alexa680-BMPP (left) or non-reactive, hydrolyzed Alexa Fluor 680 (Alexa Fluor 680 Cont.) (right) (0.075 × 3^-3^ to 0.075 μM). X-axis of each graph represents Alexa680 intensity. (D) Cell imaging of Alexa680-BMPP-treated HeLa cells. Cells were treated with Alexa680-BMPP (left) or Alexa Fluor 680 Cont.(right) (0.025 and 0.075 μM). Compound names are indicated at the top of each panel. Compound concentrations are indicated in the Figure. Scale bar, 50 μm. (E) Enlarged images of fluorescein-BMPP- (upper left panel) or Alexa680-BMPP- (lower left panel) treated cells. Corresponding control compounds are shown at right. Compound names and concentrations are indicated at the top of each panel. Scale bar, 50 μm. Note that images were obtained from different regions from and (D).

**Figure 2.**
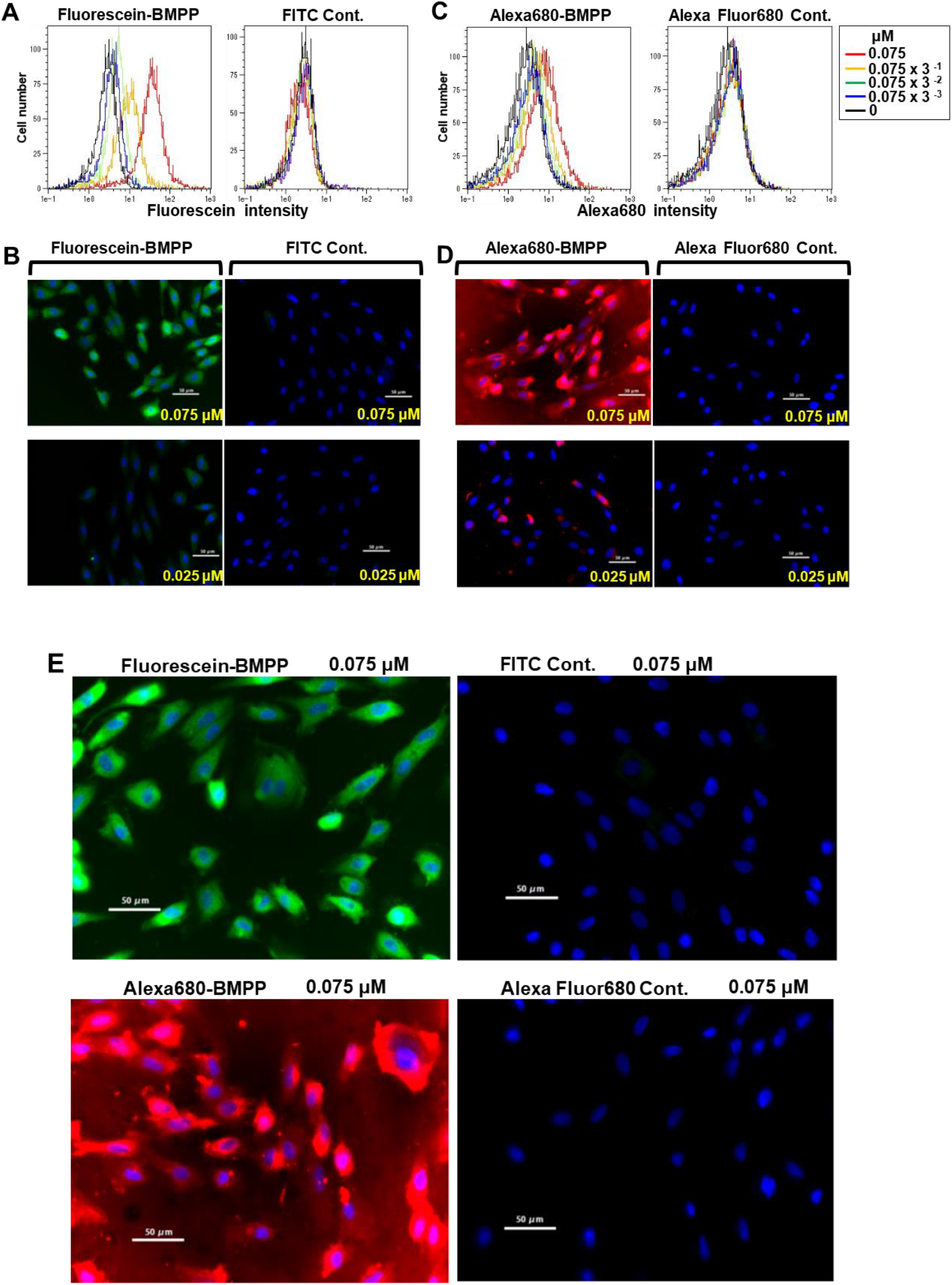
Dose-dependent import of fluorescent BMPP into rat H9C2 cells. (A) FACS analysis of fluorescein-BMPP-treated H9C2 cells. (B) Cell imaging of fluorescein-BMPP-treated H9C2 cells. Scale bar, 50 μm. (C) FACS analysis of Alexa680-BMPP-treated H9C2 cells. (D) Cell imaging of Alexa680-BMPP-treated H9C2 cells. Scale bar, 50 μm. (E) Enlarged images of fluorescein-BMPP-(upper left panel) or Alexa680-BMPP (lower left panel) treated cells. Scale bar, 50 μm. Figure is displayed using the same format as described in the legend of Figure 1.

### 3.2 Fluorescein-BMPP was imported into HeLa cells via a CD36-dependent LCFA transport pathway

We assumed that fluorescent BMPP could be imported into cells in the same manner as ^123^I-BMIPP, in which the CD36 transporter is used, because the uptake of ^123^I-BMIPP into cardiomyocytes is known to occur via CD36, a well-known LCFA transporter, reflecting the dynamics of LCFA [11-13]. Therefore, we investigated the CD36-dependent import of fluorescein-BMPP into the cells using a CD36 inhibitor. FACS analysis and cell imaging showed that Sulfosuccinimidyl Oleate (SSO), an inhibitor of CD36 [14], inhibited the import of fluorescein-BMPP into HeLa cells in a dose-dependent manner (Fig.3), strongly suggesting that fluorescent BMPP is imported into HeLa cells via CD36. Notably, even though cells were observed under 375 μM SSO conditions by cell imaging, the collected cells were largely lost in FACS analysis. This was likely due to the damaged cells under higher SSO concentration. In contrast, SSO did not inhibit H9C2 cells regarding fluorescein-BMPP import activity (data not shown). H9C2 cells appear to possess other pathways that compensate for CD36-dependent LCFA import, although the precise mechanisms remain unclear.

**Figure 3.**
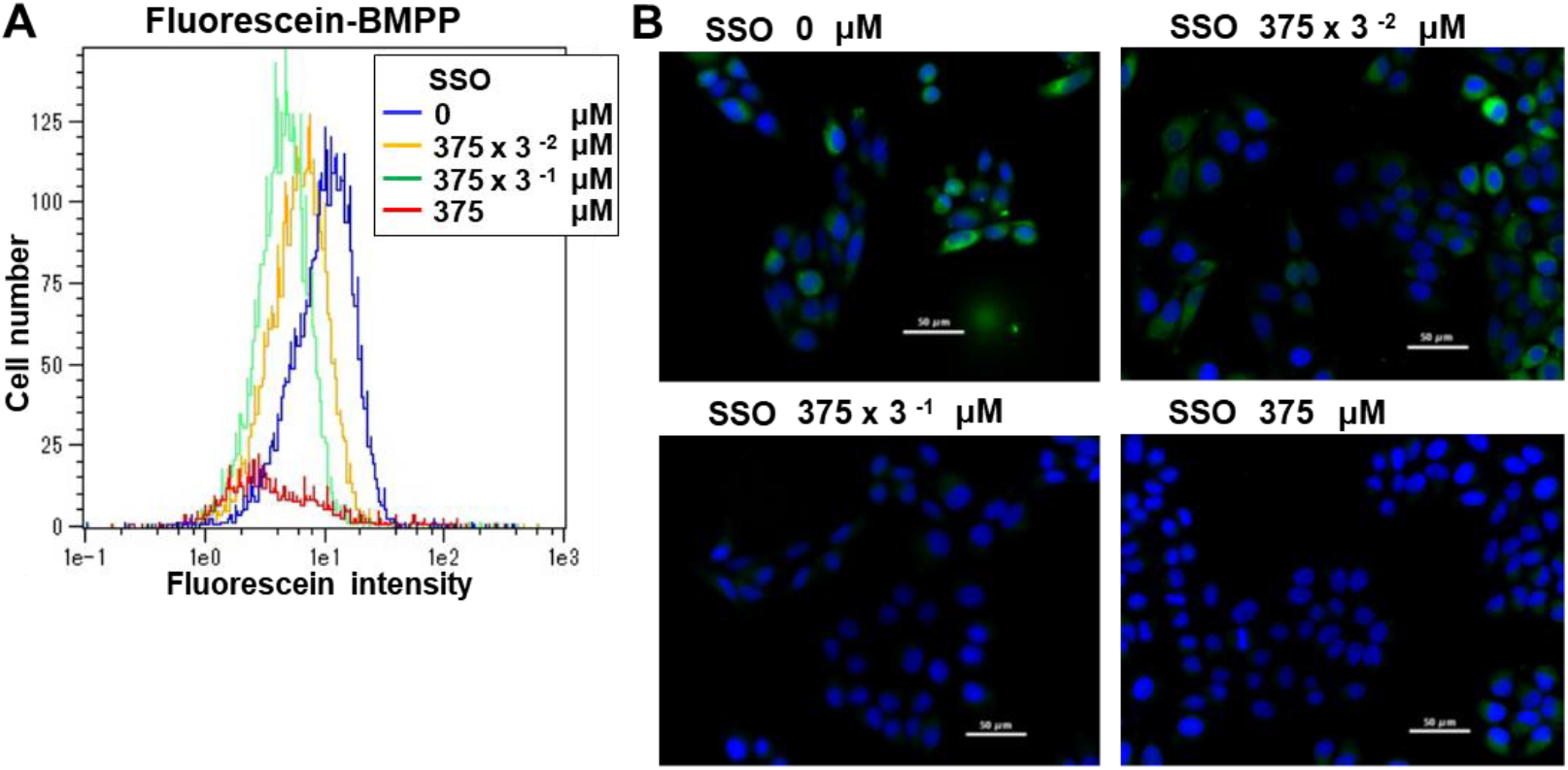
Inhibition of import of fluorescein-BMPP into HeLa cells by a CD36 inhibitor, SSO. HeLa cells were pre-treated with increasing concentrations (375 × 3 ^-2^ to 375 μM) of SSO for 30 min and co-treated for a further 30 min with both 0.05 μM fluorescein-BMPP and the same concentrations of SSO as pre-treatment. Then, fluorescein-BMPP-import analysis was performed. (A) FACS analysis of fluorescein-BMPP import into the cells. SSO concentrations are shown in the graph. (B) Cell imaging of fluorescein-BMPP import into the cells. The SSO concentrations are shown in panels. Scale bar, 50 μm. The same concentration of vehicle (DMSO) was used as 0 μM SSO.

### 3.3 Fluorescein-BMPP was exported from cells following its import into cells

To develop a fluorescently-labeled BMPP as an LCFA probe for the study of the LCFA metabolism of cells, we evaluated whether the attenuation of fluorescein-BMPP intensity in cells was detected by FACS analysis and cell imaging following import. The cells were treated with fluorescein-BMPP and then subjected to either FACS analysis immediately after treatment or cultured in non-fluorescein-BMPP medium prior to FACS analysis. Immediately after fluorescein-BMPP treatment, the cells showed an increased fluorescein intensity, in line with the results described above (Fig. 4A and C). When cells were cultured in non-fluorescein-BMPP medium following fluorescein-BMPP treatment, Alexa680 intensity was clearly attenuated in a time-dependent manner in both HeLa and H9C2 cells (Fig. 4A and C). Next, fluorescein-BMPP-treated cells were fluorescently imaged either immediately after treatment or after culturing in non-fluorescein-BMPP medium for the indicated time periods. The fluorescein-BMPP intensity was attenuated in a time-dependent manner in both HeLa and H9C2 cells (Fig. 4B and D). This attenuation of fluorescein-BMPP in cells occurring in a relatively short time (1.5 to 5 h) is in line with the fact that the wash out of ^123^I-BMIPP from myocardium is sufficiently observed in under 4 h in mice [15] and humans [16,17]. Thus, these results indicate that Alexa680-BMPP, which is unaffected by a fluorescent compound, can be exported from cells in a manner similar to ^123^I-BMIPP.

**Figure 4.**
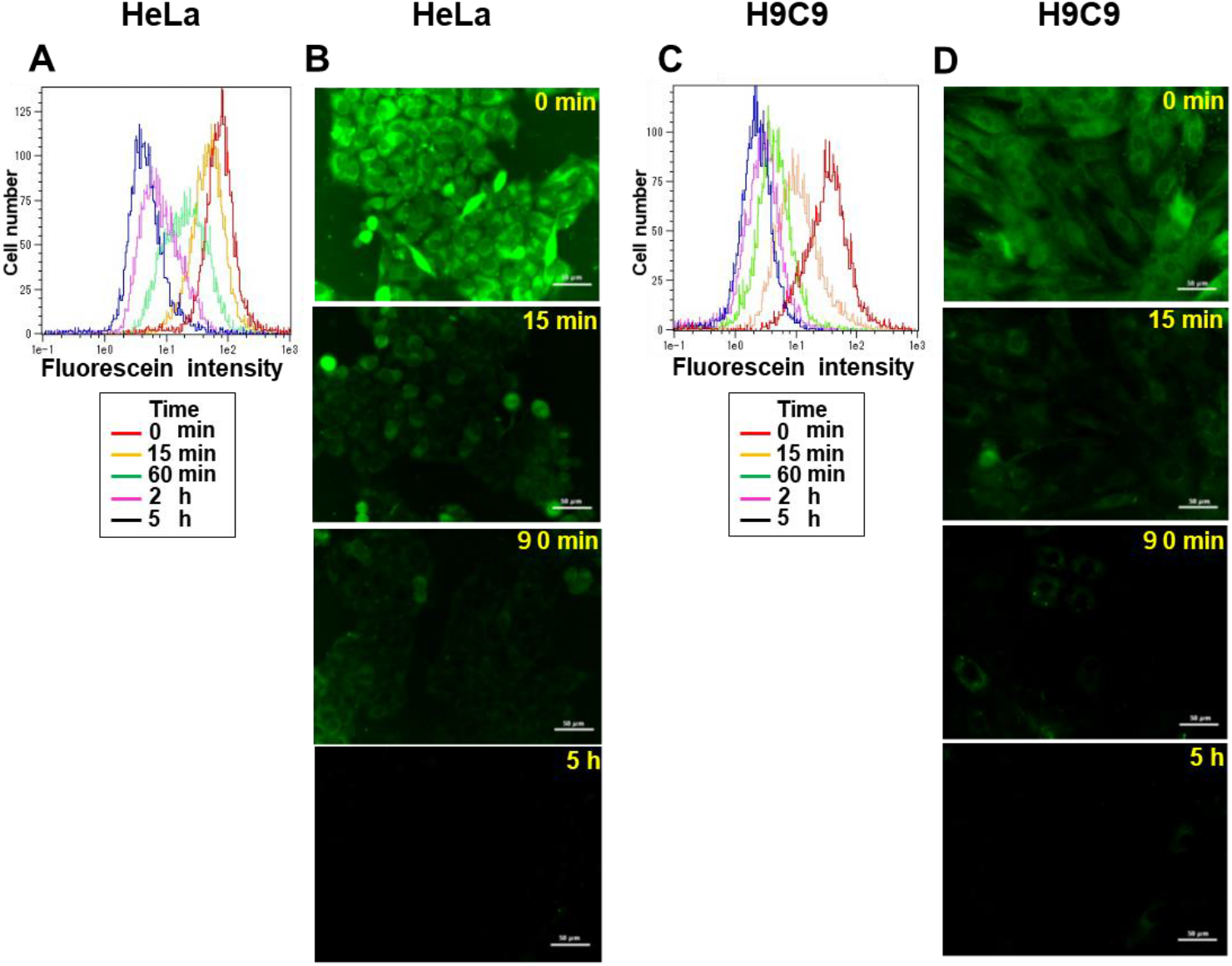
Time dependent Export of fluorescein-BMPP from cells. Cells were treated with fluorescein-BMPP (0.075 μM) for 30 min at 37 °C. The cells for measurement immediately after import (0 min) were recovered and analyzed using FACS and cell imaging. On the other hand, cells for measuring the export, in which medium was replaced to non-fluorescein-BMPP medium following the treatment, were incubated at 37 °C for time periods indicated in the figure (15 min to 5 h), and then analyzed by FACS and cell imaging. (A) FACS analysis of HeLa cell export. (B) Imaging of HeLa cell export. Scale bar, 50 μm. (C) FACS analysis of H9C2 cell export. (D) Imaging of H9C2 cell export. The cell names are indicated at the top of each panel. Scale bar, 50 μm. Hoechst 33342 staining was not performed.

## 4. Discussion

Given that ^123^I-BMIPP is commonly used as an LCFA-mimicking tracer in nuclear cardiology, we aimed to develop a potent LCFA probe with the basic structure of BMIPP for the *ex vivo* investigation of intracellular LCFA metabolism. The most distinctive feature of ^123^I-BMIPP is its methyl group at the β-3 position, which inhibits direct mitochondrial β-oxidation and is retained in cardiomyocytes for a relatively long period of time [18,19]. This feature allows ^123^I-BMIPP to be a useful tracer for comparing lipase-dependent TG metabolism in cardiomyocytes. We previously performed myocardial scintigraphy with ^123^I-BMIPP in patients with TGCV, which showed an excessive accumulation of TG in the myocardium, and scintigraphy successfully detected prolonged retention of ^123^I-BMIPP in the heart. Therefore, it has been established as a test for TGCV [3-5]. In a previous study, while utilizing the advantages of ^123^I-BMIPP, we developed an Alexa680-labeled BMIPP skeleton [10]. In the present study, we focused on the import and export ability of fluorescently-labeled BMIPP skeletons (fluorescent BMPP) in cultured cells. To the best of our knowledge, this is the first study to use a BMIPP-specific skeleton to examine the LCFA export ability of cultured cells using FACS analysis and cell imaging. An advantage of this system that uses fluorescent BMPP-and-cultured cells is that specific apparatuses are not required owing to non-radioactive detection; therefore, this method can be applied to various cells in research laboratories.

Using human HeLa cells and H9C2 cells derived from rat cardiomyocytes, we confirmed that both cell types had the ability to import fluorescent BMPP. We also confirmed that SSO, a CD36 inhibitor, inhibits the import ability of HeLa cells. CD36 is known to play a critical role in the import of ^123^I-BMIPP into cardiomyocytes [20,21]. To date, several studies have reported that the uptake of ^123^I-BMIPP into the heart is significantly impaired in CD36-deficient humans and CD36-knockout mice, suggesting that CD36 is central to the uptake of ^123^I-BMIPP into the myocardium [12,22,23]. The result of the SSO inhibition clearly indicated that fluorescent BMPP was imported into HeLa cells in a CD36-dependent manner, which is in good agreement with cardiomyocytes *in vivo*. In contrast, H9C2 cells did not show an inhibition of import activity by SSO, even though H9C2 cells were derived from rat cardiomyocytes. This result is in agreement with a previous report that SSO did not affect palmitate uptake by H9c2 cells; therefore, palmitate uptake may be mediated by CD36-independent mechanisms in H9c2 cells [24].

This study has some limitations. Fluorescent BMPP was successfully exported from cultured cells by FACS analysis and cell imaging. We believe that this export can be attributed to LCFA-metabolism-mediated transport. However, the molecular mechanism of fluorescent BMPP-export via LCFA metabolism was not directly examined in this study.

The reasons for the incorporation of fluorescent BMPP into the LCFA metabolism pathway are as follows. First, fluorescent BMPP is imported into cells depending on its BMIPP skeleton, which has been shown to be incorporated into the TG pool in cardiomyocytes *in vivo* [25]. Second, fluorescent BMPP has been reported to be imported into and exported from the hearts of mice depending on the BMIPP skeleton *in vivo* [10]. Third, ^123^I-BMIPP has been shown to be useful for examining LCFA flux under various heart conditions in humans [26,27]. Hence, Alexa680-BMPP, which has a BMIPP skeleton, is likely to enter the same metabolism as ^123^I-BMIPP once it is successfully imported into cells. However, the export of fluorescent BMPP from cells in the LCFA metabolic pathway should be elucidated in future studies.

In conclusion, this study presents a novel fluorescently-labeled LCFA probe with a basic structure of BMIPP, which allows the detection of LCFA import ability and metabolism in cultured cells, using FACS analysis and fluorescent cell imaging. The import of the probe into cells occurred, at least partially, via the LCFA transporter CD36. The probe was exported from cells following its import. Thus, this probe has great potential for the study of LCFA metabolism in a variety of cultured cells.

## Authors’ contributions

YH designed the experimental protocol, performed experiments, and wrote the manuscript. KH designed the study and coordination and wrote the manuscript. The authors have read and approved the final manuscript.

## Acknowledgements

We are grateful to Professors Kenji Monde (Hokkaido University) and Takashi Jin (Center for Biosystems Dynamics Research, RIKEN) for providing fluorescently-labeled BMPP. We thank all participants.

This study was partially supported by research grants (Translational Research Program; Strategic promotion for practical application of innovative medical technology (TR-SPRINT)) from the Japan Agency of Medical Research and Development (AMED) (Grant No. 20lm0203007j0004).

## References

[1] K. Hirano, Y. Ikeda, N. Zaima, Y. Sakata, G. Matsumiya, Triglyceride deposit cardiomyovasculopathy, N Engl J Med 359 (2008) 2396–2398. 10.1056/NEJMc0805305.

[2] K. Hirano, T. Tanaka, Y. Ikeda, S. Yamaguchi, N. Zaima, K. Kobayashi, A. Suzuki, Y. Sakata, Y. Sakata, K. Kobayashi, T. Toda, N. Fukushima, H. Ishibashi-Ueda, D. Tavian, H. Nagasaka, S.P. Hui, H. Chiba, Y. Sawa, M. Hori, Genetic mutations in adipose triglyceride lipase and myocardial up-regulation of peroxisome proliferated activated receptor-gamma in patients with triglyceride deposit cardiomyovasculopathy, Biochem Biophys Res Commun 443 (2014) 574–579. 10.1016/j.bbrc.2013.12.003.

[3] M. Li, K. I. Hirano, Y. Ikeda, M. Higashi, C. Hashimoto, B. Zhang, J. Kozawa, K. Sugimura, H. Miyauchi, A. Suzuki, Y. Hara, A. Takagi, Y. Ikeda, K. Kobayashi, Y. Futsukaichi, N. Zaima, S. Yamaguchi, R. Shrestha, H. Nakamura, K. Kawaguchi, E. Sai, S. P. Hui, Y. Nakano, A. Sawamura, T. Inaba, Y. Sakata, Y. Yasui, Y. Nagasawa, S. Kinugawa, K. Shimada, S. Yamada, H. Hao, D. Nakatani, T. Ide, T. Amano, H. Naito, H. Nagasaka, K. Kobayashi, T. s. g. Japan, Triglyceride deposit cardiomyovasculopathy: a rare cardiovascular disorder, Orphanet J Rare Dis 14 (2019) 134. 10.1186/s13023-019-1087-4.

[4] K. Hirano, Y. Ikeda, K. Sugimura, Y. Sakata, Cardiomyocyte steatosis and defective washout of iodine-123-beta-methyl iodophenyl-pentadecanoic acid in genetic deficiency of adipose triglyceride lipase, Eur Heart J 36 (2015) 580. 10.1093/eurheartj/ehu417.

[5] C. Aoshima, S. Fujimoto, A. Kudo, Y.O. Kawaguchi, K. Takamura, Y. Matsue, T. Kato, Y. Kawamura, S. Kimura, Y. Kamo, Y.O. Nozaki, D. Takahashi, N. Tomizawa, M. Hiki, T. Kasai, S. Nojiri, H. Miyauchi, K.I. Hirano, K. Shimada, K. Murakami, T. Minamino, Clinical significance of (123)I-BMIPP washout rate in patients with uncertain chronic heart failure, Eur J Nucl Med Mol Imaging 49 (2022) 3129–3139. 10.1007/s00259-022-05749-1.

[6] N. Isobe, T. Toyama, H. Hoshizaki, S. Oshima, K. Taniguchi, [Usefulness of 201Tl/123I-BMIPP myocardial SPECT to evaluate myocardial viability and area at risk in acute myocardial infarction: comparison with 201Tl/99mTc-PYP dual SPECT], Kaku Igaku 34 (1997) 213–220.

[7] T. Mochizuki, K. Murase, H. Higashino, M. Miyagawa, Y. Sugawara, T. Kikuchi, J. Ikezoe, Ischemic “memory image” in acute myocardial infarction of 123I-BMIPP after reperfusion therapy: a comparison with 99mTc-pyrophosphate and 201Tl dual-isotope SPECT, Ann Nucl Med 16 (2002) 563–568. 10.1007/BF02988634.

[8] S. Matsuo, K. Nakajima, S. Kinuya, M. Yamagishi, Diagnostic utility of 123I-BMIPP imaging in patients with Takotsubo cardiomyopathy, J Cardiol 64 (2014) 49–56. 10.1016/j.jjcc.2013.10.019.

[9] S.K. Biswas, M. Sarai, H. Hishida, Y. Ozaki, 123I-BMIPP fatty acid analogue imaging is a novel diagnostic and prognostic approach following acute myocardial infarction, Singapore Med J 50 (2009) 943–948.

[10] M.M.M. Swamy, M.Z.M. Zubir, Mutmainah, S. Tsuboi, Y. Murai, K. Monde, K.I. Hirano, T. Jin, A near-infrared fluorescent long-chain fatty acid toward optical imaging of cardiac metabolism in living mice, Analyst 147 (2022) 4206–4212. 10.1039/d2an00999d.

[11] M. Yuasa-Kawase, D. Masuda, T. Yamashita, R. Kawase, H. Nakaoka, M. Inagaki, K. Nakatani, K. Tsubakio-Yamamoto, T. Ohama, A. Matsuyama, M. Nishida, M. Ishigami, T. Kawamoto, I. Komuro, S. Yamashita, Patients with CD36 deficiency are associated with enhanced atherosclerotic cardiovascular diseases, J Atheroscler Thromb 19 (2012) 263–275. 10.5551/jat.10603.

[12] T. Yoshizumi, S. Nozaki, K. Fukuchi, K. Yamasaki, T. Fukuchi, T. Maruyama, Y. Tomiyama, S. Yamashita, T. Nishimura, Y. Matsuzawa, Pharmacokinetics and metabolism of 123I-BMIPP fatty acid analog in healthy and CD36-deficient subjects, J Nucl Med 41 (2000) 1134–1138.

[13] E.H. Hwang, J. Taki, S. Yasue, M. Fujimoto, M. Taniguchi, I. Matsunari, K. Nakajima, S. Shiobara, T. Ikeda, N. Tonami, Absent myocardial iodine-123-BMIPP uptake and platelet/monocyte CD36 deficiency, J Nucl Med 39 (1998) 1681–1684.

[14] O. Kuda, T.A. Pietka, Z. Demianova, E. Kudova, J. Cvacka, J. Kopecky, N.A. Abumrad, Sulfo-N-succinimidyl oleate (SSO) inhibits fatty acid uptake and signaling for intracellular calcium via binding CD36 lysine 164: SSO also inhibits oxidized low density lipoprotein uptake by macrophages, J Biol Chem 288 (2013) 15547–15555. 10.1074/jbc.M113.473298.

[15] A. Suzuki, S. Yamaguchi, M. Li, Y. Hara, H. Miyauchi, Y. Ikeda, B. Zhang, M. Higashi, Y. Ikeda, A. Takagi, H. Nagasaka, K. Kobayashi, Y. Magata, T. Aoyama, K.I. Hirano, Tricaprin Rescues Myocardial Abnormality in a Mouse Model of Triglyceride Deposit Cardiomyovasculopathy, J Oleo Sci 67 (2018) 983–989. 10.5650/jos.ess18037.

[16] Z. Chen, K. Nakajima, K.I. Hirano, T. Kamiya, S. Yoshida, S. Saito, S. Kinuya, Methods of calculating (123)I-beta-methyl-P-iodophenyl-pentadecanoic acid washout rates in triglyceride deposit cardiomyovasculopathy, Ann Nucl Med 36 (2022) 986–997. 10.1007/s12149-022-01787-9.

[17] I. Matsunari, T. Saga, J. Taki, Y. Akashi, J. Hirai, T. Wakasugi, T. Aoyama, M. Matoba, K. Ichiyanagi, K. Hisada, Kinetics of iodine-123-BMIPP in patients with prior myocardial infarction: assessment with dynamic rest and stress images compared with stress thallium-201 SPECT, J Nucl Med 35 (1994) 1279–1285.

[18] F.F. Knapp, Jr., J. Kropp, BMIPP design and development, Int J Card Imaging 15 (1999) 1–9. 10.1023/a:1006147228352.

[19] Y. Yamamichi, H. Kusuoka, K. Morishita, Y. Shirakami, M. Kurami, K. Okano, O. Itoh, T. Nishimura, Metabolism of iodine-123-BMIPP in perfused rat hearts, J Nucl Med 36 (1995) 1043–1050.

[20] J.F.C. Glatz, J. Luiken, Dynamic role of the transmembrane glycoprotein CD36 (SR-B2) in cellular fatty acid uptake and utilization, J Lipid Res. 59 (2018) 1084–1093. 10.1194/jlr.R082933.

[21] N.A. Abumrad, I.J. Goldberg, CD36 actions in the heart: Lipids, calcium, inflammation, repair and more?, Biochim Biophys Acta 1861 (2016) 1442–1449. 10.1016/j.bbalip.2016.03.015.

[22] C.T. Coburn, F.F. Knapp, Jr., M. Febbraio, A.L. Beets, R.L. Silverstein, N.A. Abumrad, Defective uptake and utilization of long chain fatty acids in muscle and adipose tissues of CD36 knockout mice, J Biol Chem 275 (2000) 32523–32529. 10.1074/jbc.M003826200.

[23] T. Tanaka, T. Nakata, T. Oka, T. Ogawa, F. Okamoto, Y. Kusaka, K. Sohmiya, K. Shimamoto, K. Itakura, Defect in human myocardial long-chain fatty acid uptake is caused by FAT/CD36 mutations, J Lipid Res 42 (2001) 751–759.

[24] F.A. Van Nieuwenhoven, J.J. Luiken, Y.F. De Jong, P.A. Grimaldi, G.J. Van der Vusse, J.F. Glatz, Stable transfection of fatty acid translocase (CD36) in a rat heart muscle cell line (H9c2), J Lipid Res 39 (1998) 2039–2047.

[25] S. Morishita, H. Kusuoka, Y. Yamamichi, N. Suzuki, M. Kurami, T. Nishimura, Kinetics of radioiodinated species in subcellular fractions from rat hearts following administration of iodine-123-labelled 15-(p-iodophenyl)-3-(R,S)-methylpentadecanoic acid (123I-BMIPP), Eur J Nucl Med 23 (1996) 383–389. 10.1007/BF01247365.

[26] H. Seki, T. Toyama, K. Higuchi, S. Kasama, T. Ueda, R. Seki, T. Hatori, K. Endo, M. Kurabayashi, Prediction of functional improvement of ischemic myocardium with (123I-BMIPP SPECT and 99mTc-tetrofosmin SPECT imaging: a study of patients with large acute myocardial infarction and receiving revascularization therapy, Circ J 69 (2005) 311–319. 10.1253/circj.69.311.

[27] A.S. Hambye, A.A. Dobbeleir, A.M. Vervaet, P.A. Van den Heuvel, P.R. Franken, BMIPP imaging to improve the value of sestamibi scintigraphy for predicting functional outcome in severe chronic ischemic left ventricular dysfunction, J Nucl Med 40 (1999) 1468–1476.

